# CyNET- a network analysis framework for high dimensionality, system level analyses of the functional Immunome

**DOI:** 10.1101/2024.07.15.603495

**Authors:** Pavanish Kumar, Joo Guan Yeo, Su Li Poh, Sharifah Nur Hazirah, Nursyuhadah Binte Sutamam, WX Gladys Ang, Vasuki Ranjani Chellamuthu, Martin Wasser, Yeo Kee Thai, Jing Yao Leong, Lakshmi Ramakrishna, Thaschawee Arkachaisri, Salvatore Albani

## Abstract

The immune system is a complex “Network of Networks”, in which various immune cell subsets interact and influence each other’s functions, ultimately determining immune competence and the control or onset of disease. The coordinated interactions between these cell subsets determine whether the outcome is a normal physiological state or a pathological condition. Established statistical procedures largely ignore the interactions between subsets and rely on statistically significant changes in the frequency of cell subsets. We developed CyNET (**Cy**tometry **Net**work)— an analysis platform based on a network science approach to understand the immune system holistically. CyNET enables us to quantify the systems level and subsets level properties. We show that changes in the centrality of the nodes (immune subsets) reflect better biological functions than changes in frequency. We used CyNET to analyze the immune development along the chronological age gradient. Peripheral blood cells from healthy newborns (cord blood), adults (20 to 55 years), and elderly (70 years and above) human subjects were further used for validation using single-cell transcriptomics. We found that network edge density, degree centralization, and assortativity score reflect the maturation and development of the immune system along the age axis, thus enabling the characterisation of the functional architecture of the Immunome and the identification of key functional hubs within the immune networks.

## Introduction

High dimensionality approaches, in particular single cell proteomics using Mass Cytometry (CyTOF) and multiparametric fluorescent activated cell sorting (FACS) to include a larger number of markers available for staining, have contributed immensely towards our understanding of the immunome as a network of networks. High-dimensional cytometry has its challenges and requires innovative data- driven machine-learning (ML) approaches to understand the data generated. For example, a typical flow cytometry analysis requires biaxial plots and existing knowledge/hypothesis-driven sequential gating to characterize cell types. A test with just 30 markers requires 870 bi-axial plots and one with 40 markers will require 1560 bi-axial plots. It is therefore very challenging or probably impossible for humans to conceive and visualize over 1000 biaxial plots together to make sense of it, thus preventing the discovery of unexpected combinations of protein expression on a cell type and, most importantly, limiting the ability to understand the true architecture of the immunome, and the relevance of interactions among functionally relevant subsets. This need inspired an interdisciplinary collaboration between data scientists, ML scientists, and immunologists, thereby sparking a novel way of looking at these complex cytometric data. ML methods in concert with immunology expertise can lead to the development of approaches, that allow unbiased characterization and discovery of immune cell types and functions. Newer visualization methods (1–3) allowed a holistic understanding of the immune system. These developments are particularly relevant for clinical and translational research, and have sparked new approaches for data-driven theragnostics tools for precision medicine (4–6) and clinical development (7,8)

However, approaches that consider the relationship among the components of the immune response at system levels are still lacking. In real *in-vivo* conditions, various immune cell subsets interact and influence each other’s functions. The coordinated interactions between these immune cell subsets eventually determine the outcome, reflected as a normal physiological state or pathological condition. Established statistical procedures used in cytometry analyses ignore the interactions between subsets and rely entirely on statistically significant changes in the frequency or percentage of cell subsets.

Also, no previous method has explored the understanding of the Immunome at a systems level yet, and no measures have been investigated and developed to understand the properties of the system. We have explored the core idea of Systems Immunomics (SI) in previous work while studying an array of human diseases (9,8,10,11). In the present study, we developed the core idea further and created a web application software named CyNET to facilitate the availability of an SI approach to the broader community of immunologists and clinician scientists. We tested our approach by analysing the immunome of healthy individuals from newborn (cord blood), adult (20 to 55 years), and elderly (70 years and above) human subjects using CyNET. We further performed single-cell transcriptomics analysis to validate the findings from CyNET network analysis. We propose here the concept that centrality of given immune cell subsets within the network may be functionally relevant, and integrates the use of frequency as the only parameter employed for a comprehensive understanding of the Immunone in health and disease.

## Results

### Network analysis framework

To understand the healthy immunome, we analyzed peripheral blood mononuclear cells (PBMCs) using CyTOF. The data was obtained from the EPIC study (5). The samples from CBNB (newborn cord blood, n=17), adult (25-55 years, n=17), and elderly human subjects (70 years and above, n=17) were analyzed using CyNET software (see Methods). The analyzed data included PBMCs stained with two CyTOF panels. Panel 1 (P1) was focused on T and NK cells, utilising PMA/ionomycin stimulation to facilitate the identification of chemokine and cytokine-secreting cells, while Panel 2 (P2) was focused on detection of unstimulated monocytes/DCs and B cells. P1 and P2 data for the 3 age groups were analyzed using the EPIC analytics platform. Briefly, data was batch-normalized and clustered into 100 subsets. The frequencies of the subsets were calculated for correlation network analysis using CyNET. Subset phenotypes were identified using median marker intensity (Fig.S1). The frequency of T and NK cell subsets from P1, B cells, and monocytes/DC subsets from P2 were then used to build the immune cell network (Fig1). Immune cell clusters formed the nodes of the network and to establish edge/connection between the clusters correlation coefficients >= 0.6 and p val < 0.05 were used. The network was visualized using force-directed layout algorithms from the igraph and visNetwork R packages. Each immune cell network’s various system-level properties were calculated (Table 1). To calculate the network’s modularity, the clusters were grouped into major lineage types as shown in Fig.1. Strong correlations between the subsets reflect potential direct or indirect interaction between the subsets.

**Fig. 1.**
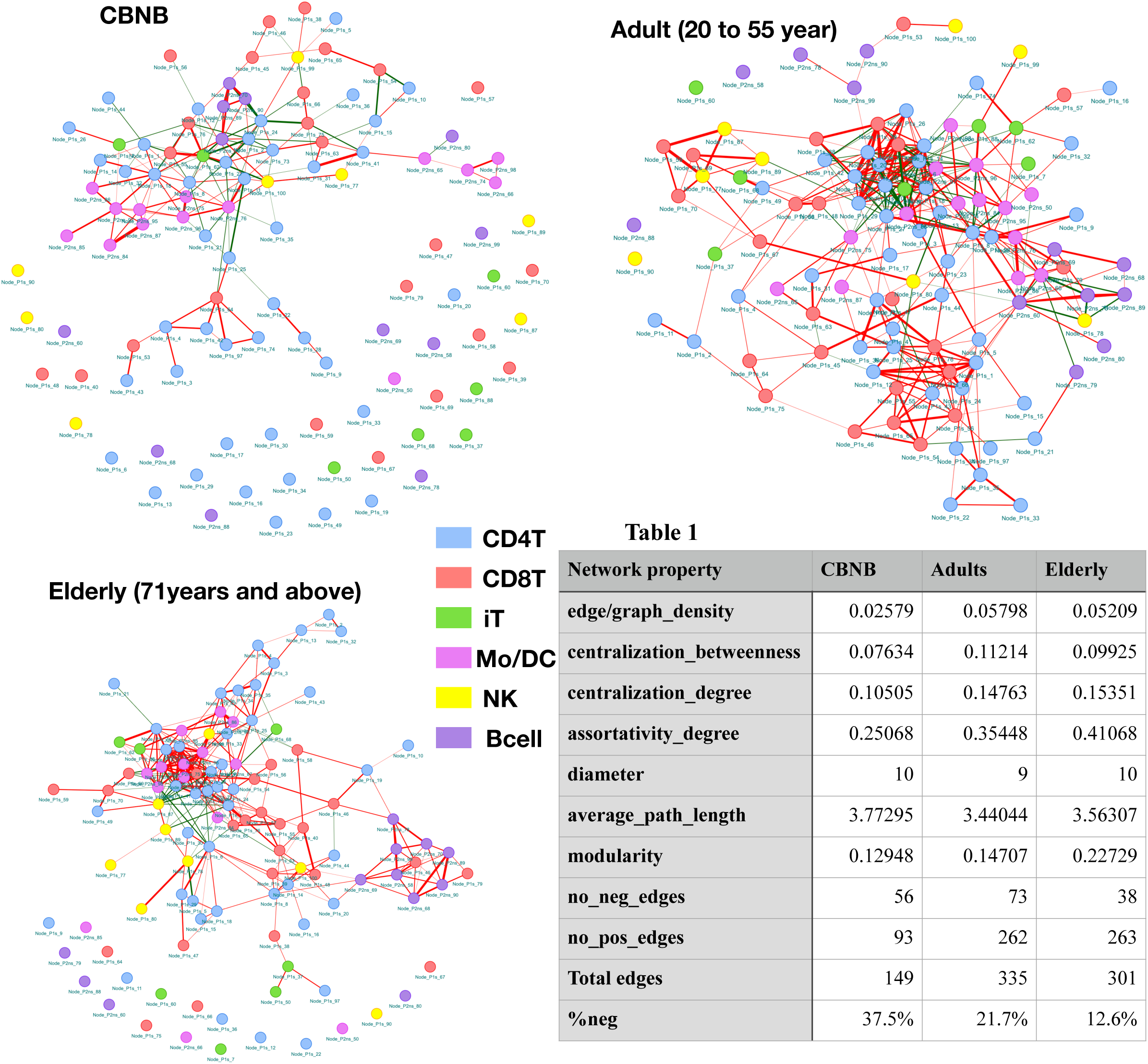
Immune cell network of CBNB, Adults and elderly. Immune cell network for each group was created where nodes are cellular subsets and edges are potential interaction between the subsets. Nodes are colour coded based on major immune cell lineage. Edges in green reflects negative correlation (inhibitory interaction) and red reflects positive interaction between the subsets. Thickness of the edges shows the strength of interaction (magnitude of correlation). A, B, C) Shows immune cell network of cord blood new born (CBNB), adults and elderly group respectively. D) Shows network property in a table format.

### Node Degree centrality reflects the changes in biological function, in some cases better than the frequency

Communication hubs within the network are often pivotal for the appropriate functionality of the entire immunome. Thus, identifying the functional hubs and mechanistic drivers and how they relate to each other within the immunome is essential to understand its functionality. Here, we used Degree of Centrality (DC) to measure and identify the communication hub and drivers of immune homeostasis. We hypothesized that nodes with higher frequency, which denote higher abundance immune subsets, are functionally more important and may have higher interactions with other subsets. To our surprise, we observed that nodes with higher frequency do not always show a higher DC. The correlation analysis between the median frequency and DC showed no significant correlation in either CBNB (r=0.099, p=0.391), adult (r= 0.151, p=0.121) or elderly (r=0.026, p=0.802) group (Fig.1A). Consequently, we systematically investigated the immune cell subset properties for the major cell lineage groups.

#### T cells

There were 43 CD4 T cell subset nodes (Fig. S2A), which accounted for maximum edges in all three networks (Fig.S3). CD4 T cell subsets formed the majority of cells in CBNB (median = 60 %) while in adults (41%) and elderly (44%) it was lower than CBNB (Fig.S4). Among the 43 CD4 T cell nodes only a few were more enriched in CBNB than in adult and elderly subjects (Fig. S2A). These more abundant CD4T cell nodes from CBNB (Nodes 1, 2,25,34,44,73) expressed CD45RA, a naive T cell marker, explaining their lower interactions and the lesser DC(Fig.2B).

**Fig. 2.**
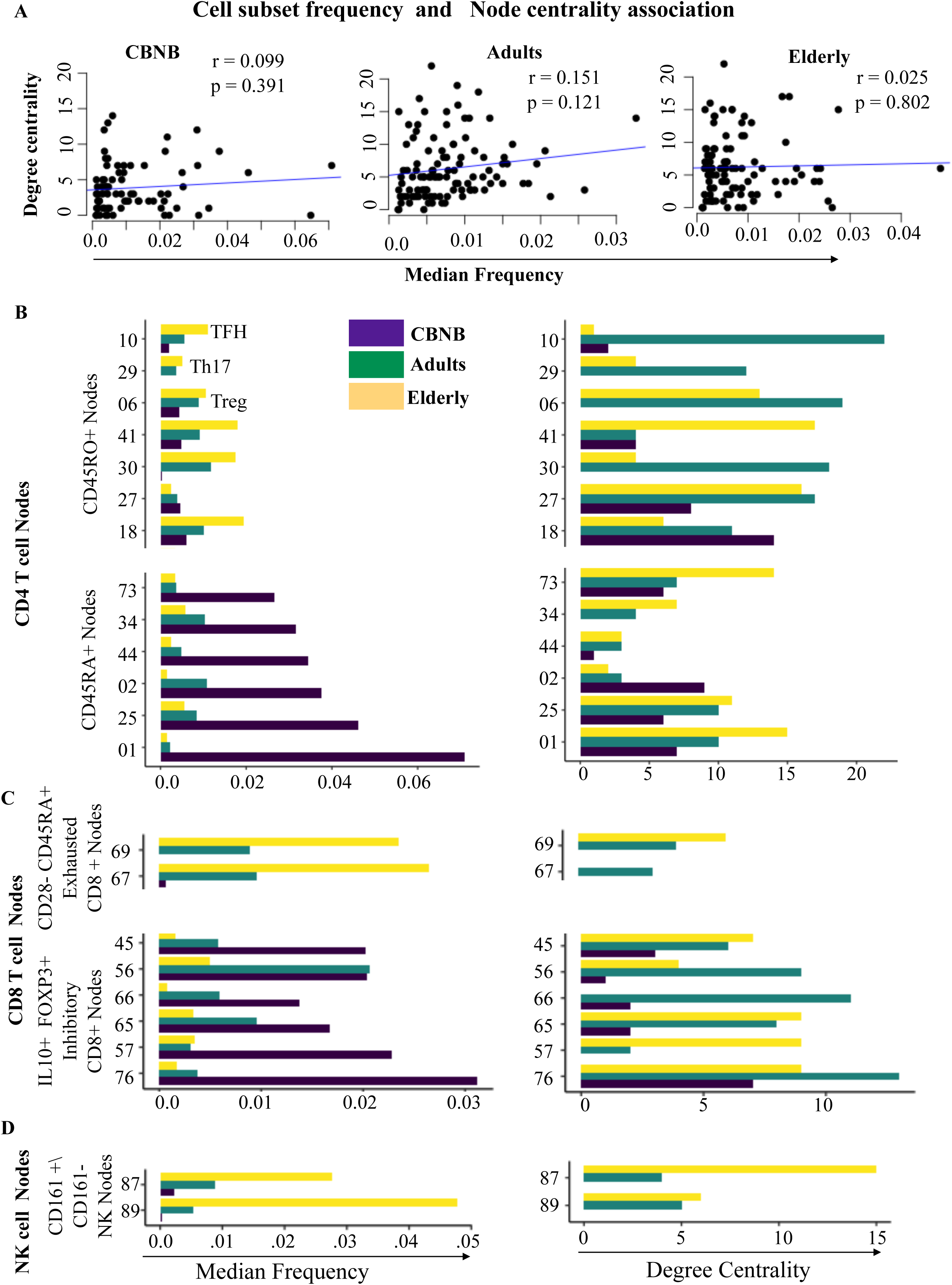
Association between Degree Centrality (DC) and frequency of immune cell subsets. A) Median frequency and DC of all immune cell subsets in CBNB, adult and elderly group was plotted as dot charts where each dot shows a node/subset. Blue line shows the linear regression line showing linear association between frequency and DC. Correlation coefficient (Pearson correlation) and P-value was also shown on each group plot. Median frequency and DC of selected node in B) CD4 T cell subsets C) CD8 T cell subset D) NK cell subsets were plotted as bar-plot. colour of the bars in bar-plot shows the human sample group information. Each node number is shown on Y -axis while median frequency and DC was shown on X-axis.

In contrast, memory CD4T cell (CD45RO+) subsets (Nodes 18, 27, 30,41,6) showed a higher DC than the CD45RA+ nodes (Fig.2A). We investigated the phenotype of the most central CD4 T cell node (Node10). Node10 showed a phenotype (CXCR5+PD1+) of Tfh (T follicular helper cells) (Fig.S1A). This Tfh subset had the highest DC (DC = 22) in the adult network while in CBNB (DC=2) and elderly (DC =1) immune cell networks its DC was much lower (Fig.2B). Similarly, a Th17 subset (Node 29) had lower frequency in all three groups compared with many other CD4 T cell nodes, however this node (Node 29) showed higher DC in adults’ network (Fig.2B). Although, CBNB samples had the highest fraction of CD4 T cells, heterogeneity was reduced, when compared to adult and elderly, where many nodes were present with similar frequency. The average DC of the CD4 T cell node in CBNB was lower (average DC = 2.35) compared to adult (average DC = 5.2) and elderly (average DC = 4.3).

Similar to CD4 T cell subsets, CD8 T cell subsets from CBNB showed a lower degree of centrality for the subsets (Node 76, 65,66,56,57,45) with higher median frequency (Fig.2C). These CBNB enriched CD8 T cell subsets expressed IL10 and FOXP3, suggesting a regulatory phenotype (12,13,14) which may play a role in tolerance induction. Furthermore, the most abundant CD8 subset (Node67) in the elderly showed 0 DC, and another abundant subset Node69 also showed lower DC (DC = 6) (Fig. 2C), conversely Node 63 showed the highest DC (DC = 15) even though its median frequency was much lower (median frequency < 0.01) (Fig. S2C). Node67 and Node69 had an exhausted phenotype (CD28- CD45RA+), and Node63, except for CD3 and CD8, was negative for most markers that we analysed (Fig.S1A).

#### Innate cells (NK, DC)

A total of 8 NK cell subsets were identified (Fig. S2C). The NK cell subsets showed lower DC in adult than in elderly subjects’ network (Fig. S2C). In CBNB, both frequency and degree of centrality were low (Fig. S2C). The most abundant NK subset (Node89) in the elderly was of a CD161+ GZMB+IFNg-low phenotype the while the subset with the highest DC (Node87) was CD161-GZMB+IFNg+ (Fig.2D, Fig.S1).

In CBNB, the most abundant monocyte subset (Node76) was HLA-DR negative and CD86 low, in contrast, the most frequent subset (Node86) in adults and the elderly was HLA-DR and CD86 positive. Here again, the DC of the most monocyte subsets in CBNB was lower compared to adult and the elderly (Fig.3A). Most abundant subsets (Node 65,66,76,96) in CBNB were negative for HLA-DR expression. HLA-DR is required for antigen presentation to T cells, and a lack of it on CBNB monocytes shows impairment or lack of cellular function by monocytes. Interestingly, the proportion of edges between monocyte subsets (12.1%) was highest in CBNB even though the CBNB network was sparse while the converse is true for adults’ immune cell network with the lowest edge fraction between the monocytes (5.4%) (Fig.S3). In the adult immune network, monocytes were connected to many subsets of other lineages, while in CBNB the monocytes were mostly connected to other monocyte subsets (Fig1).

**Fig. 3.**
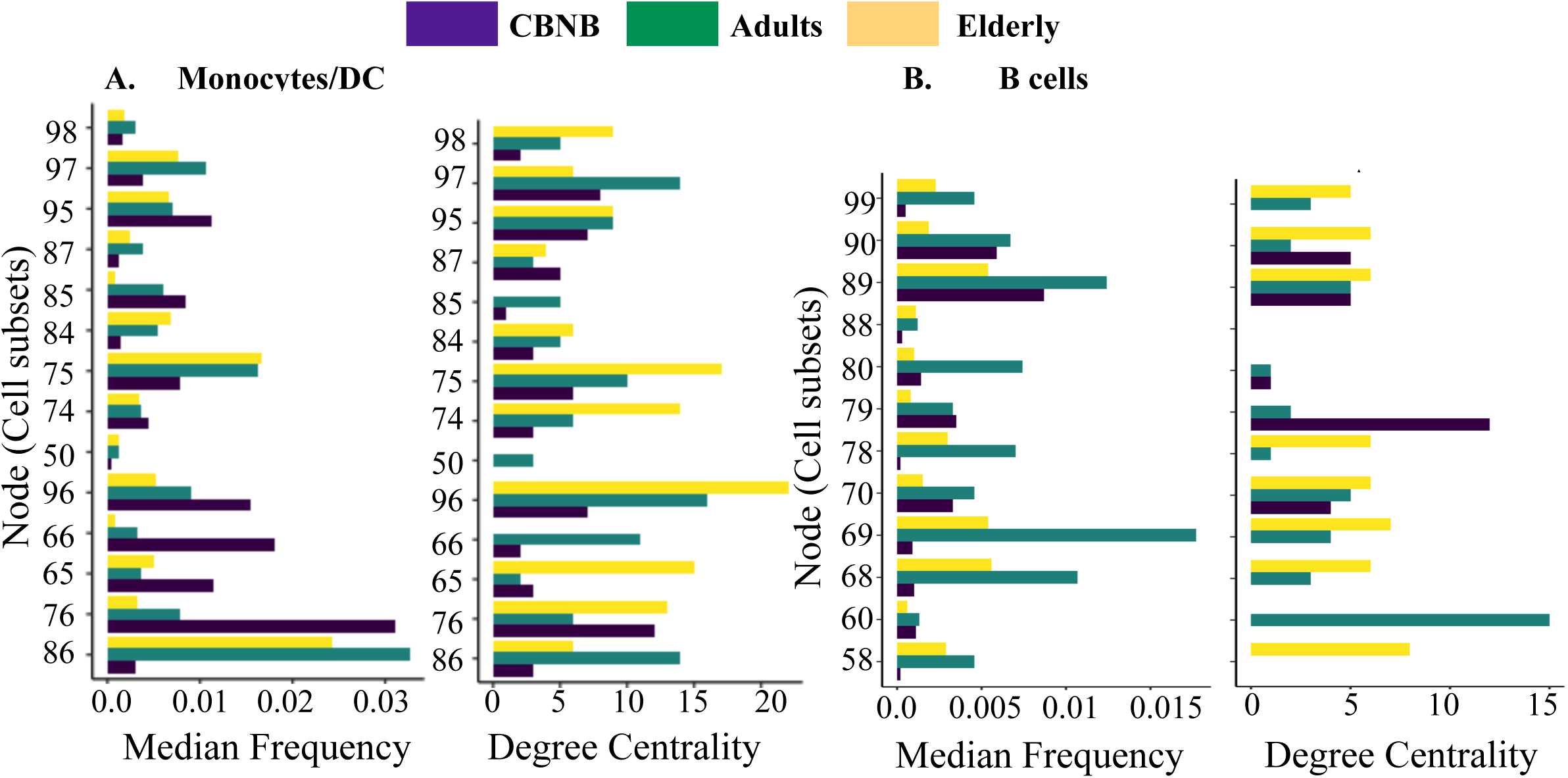
Degree Centrality (DC) and frequency of immune cell subsets from panel 2 dataset. Median frequency and DC of each node in A) Monocytes/DC cell subsets B) B cell subsets were plotted as bar-plot. colour of the bars in bar-plot shows the human sample group information. Each node number is shown on Y -axis while median frequency or DC was shown on X-axis.

#### B cells

The major function of B cells is the production of neutralizing antibodies against invading pathogens. B cell subsets were low in CBNB and elderly subjects compared to adults (Fig.S4). In CBNB, only subset (Node79) which had a phenotype (CD19+CD38+CD24+IL10+) consistent with regulatory B cells (Breg) (Fig.S1B) showed DC greater than 5 and it was not the most abundant subset in the CBNB(Fig.3B). Breg cells are involved in the early maintenance of pregnancy and thus may have functional importance in the developing fetus. In adults, a naive B cell subset (Node60) showed the highest degree centrality (Fig.3B). This subset also expressed trafficking receptor CXCR4 and low levels of plasmocytoid dendritic cell (pDC) specific receptor CD303. Both, B cells and pDC can be generated from common lymphoid precursors (CLP) and low levels of CD303 on naive cells may suggest that this subset is at earlier end of the maturation state. In the elderly immune network, B cell subsets showed edges between themselves and clustered as isolated module (Fig.1C). The isolated cluster of B cells suggests a loss of B cell cross talk/engagement in the overall immune cell network with aging.

### Network edge density, and centralization score reflect the maturation and development of the immune system along the age gradient

Tools to understand and dissect immune functions and their relations at systems level have not been developed yet. In the present study we investigated how the various elements of the Immunome interlace functionally, and hypothesise that these interactions affect the function and phenotype of individual immune functions within the network. In particular, we investigated graph density, centralisation score, assortativity score and modularity as a relevant attribute to understand and depict the development and maturation of the immune system with age.

The graph or edge density of the network is the ratio of edges observed to maximum possible edges in the network. The edge density of a network ranges from 0, for a completely isolated network, to 1 for a completely connected network, and is a measure of overall interaction in the network. We hypothesized that the developing, immature immune cell subsets in CBNB will have a lower network edge density compared to adults and elderly immune cell networks. As expected, we found that the edge density was lowest for CBNB (0.0257) compared to network densities of the adult (0.0578) and elderly (0.0521) (Table 1). Lower edge density was also visually apparent in the CBNB network, where many nodes were isolated and had no edges between them (Fig.1A). Also, we found that the total number of edges (total edges = 149) in the CBNB network was lower than the network of adult (total edges = 335) and elderly (total edges = 301) human subjects. Moreover, the percentage of negatively correlated edges was higher in CBNB (37.5%) than in adults (21.7%) and elderly (12.6%) networks. The decrease in negatively correlated edges suggests a loss of counterbalancing regulatory and inhibitory functions of the network. These data provide another perspective to the notion that aged/elderly populations are more prone to pro-inflammatory conditions than younger and adult individuals.

We further analyzed the centralization score, which measures the importance and dominance of the nodes in the network. A centralization score of 0 means all nodes in the network are equally central or equally important, while a centralization score of 1 suggests only 1 node completely dominates the network. Intuitively, we hypothesized that at the earlier stages of development, immune cell networks are less centralised as specialized functions of the cells may not have emerged or developed yet, and with development and antigen experience more nodes will take a specialised role and thus will dominate the network or subnetworks. This will be reflected as higher centralization of the network. We quantified the degree centralization score of the CBNB (0.105), adult (0.148), and elderly (0.154) immune cell network, and observed an increase in the centralization score on the age axis. We also observed a concordant pattern in graph degree assortativity, which is another measure for preferential attachment between hub or highly connected network nodes(15), with an increase in degree assortativity score on the age axis (Table 1). Thus, the development of cells and exposure to stimuli with increasing age lead to the development of more specialized immune cell subsets with an increased ability to interact with other subsets, reflected as an increase in the preferential attachment or assortativity score of the network.

We also hypothesized that age would affect the modularity of immune networks. Modularity reflects inter and intra-communication of the node members within the network, where interactions within the same community increase the modularity score while interactions between communities lower the modularity. In our immune cell networks, major cell lineages were delineated as communities for modularity score calculation. We observed that modularity increased along the age axis (Table 1), indicating loss of communication between the major cell lineage subsets with age. Within this context, B cell subsets formed an isolated cluster in the elderly immune, in comparison with the higher number of inter-subset interaction of B cells in adult immune network (Fig.1). While the total B cell edges in the adult and elderly networks were 30 and 32, respectively., 36 % (11 edges) were solely between the B cell subsets in the adult network while this was much higher 56% (18 edges) in the elderly network (Fig.S3). Similarly, intra-subset edges within the monocyte/DC subsets increased from adult (19%) to elderly (28%). Overall, the immune network of CBNB was sparse, with much lower modularity than adult and elderly.

### Single-cell Transcriptomics analysis

As Ligand and Receptor (LR) interactions between cells comprise of the most common ways of cell- cell communication, we hypothesized that interactions inferred from the correlation network could be explained by LR interaction between the cell subsets. More connected nodes, as reflected in a higher degree centrality score, would thus have higher LR interaction potential with other subsets. Thus, to validate the correlational structure and degree centrality of a subset we performed LR interaction analysis between immune cell subsets from PBMCs using single-cell RNA sequencing (scRNA-seq). We performed 10x drop-seq based scRNA-seq on PBMCs from CBNB(n=5), adult(n=6), and elderly(n=4) human subjects (Table S1). After QC and filtering 22975 cells were retained for analysis (see Methods). scRNA-seq data was batch corrected using the SCT transform method, and transformed data was used for predicting the expression of genes using the ALRA zero count preserving algorithm(16). The ALRA predicted gene count matrix was then used for cell type identification. The SingleR package was used for cell type identification using immune cell signature from MonacoImmuneData from celldex R package(17). A total of 25 cell types were identified (Fig.4, Table S2) using scRNAseq. The most abundant cell types recovered in CBNB and adults were naive CD4 T cells while in the elderly NK cells were most abundant. In our immune network analysis naive CD4 T cells have lower interaction across all age groups compared to other CD4 T cell subsets such as Th17, Tfh and Tregs (Fig.2A). In agreement with these observations, differential LR gene expression analysis between naive CD4 T cells and the rest of CD4 T cells showed lower expression of 57 LR genes and higher expression of 41 LR genes in naive CD4T cells (Table S3). The low expressed genes included the key LR genes such as chemokines and cytokines receptors (*CCR4, CCR6, CCR10, CXCR3, IL10RA, IL17RB, IL2RA, TNFRSF1B*), chemokine, cytokine (*CCL5, IL2*) and integrin (*ITGB1, ITGAV, ITGA2*) genes. Among the highly expressed LR genes were growth factors such as *FLT1*, and *IGF1R*. Inhibitory receptors such as *PECAM-1* and *BTLA* were also more expressed in naive CD4T cells. The dampening of key chemokines/cytokines, chemokine/cytokine receptors, and integrin genes explains the lower interaction of naive CD4T cells with other immune cells. The CBNB immune cell network was the most sparse with lower interactions between subsets (Fig.1A). Both CyTOF (Fig.S4) and scRNA-seq data (Fig.4) showed naive CD4 T cells as the most abundant immune cells in CBNB group.

**Fig. 4.**
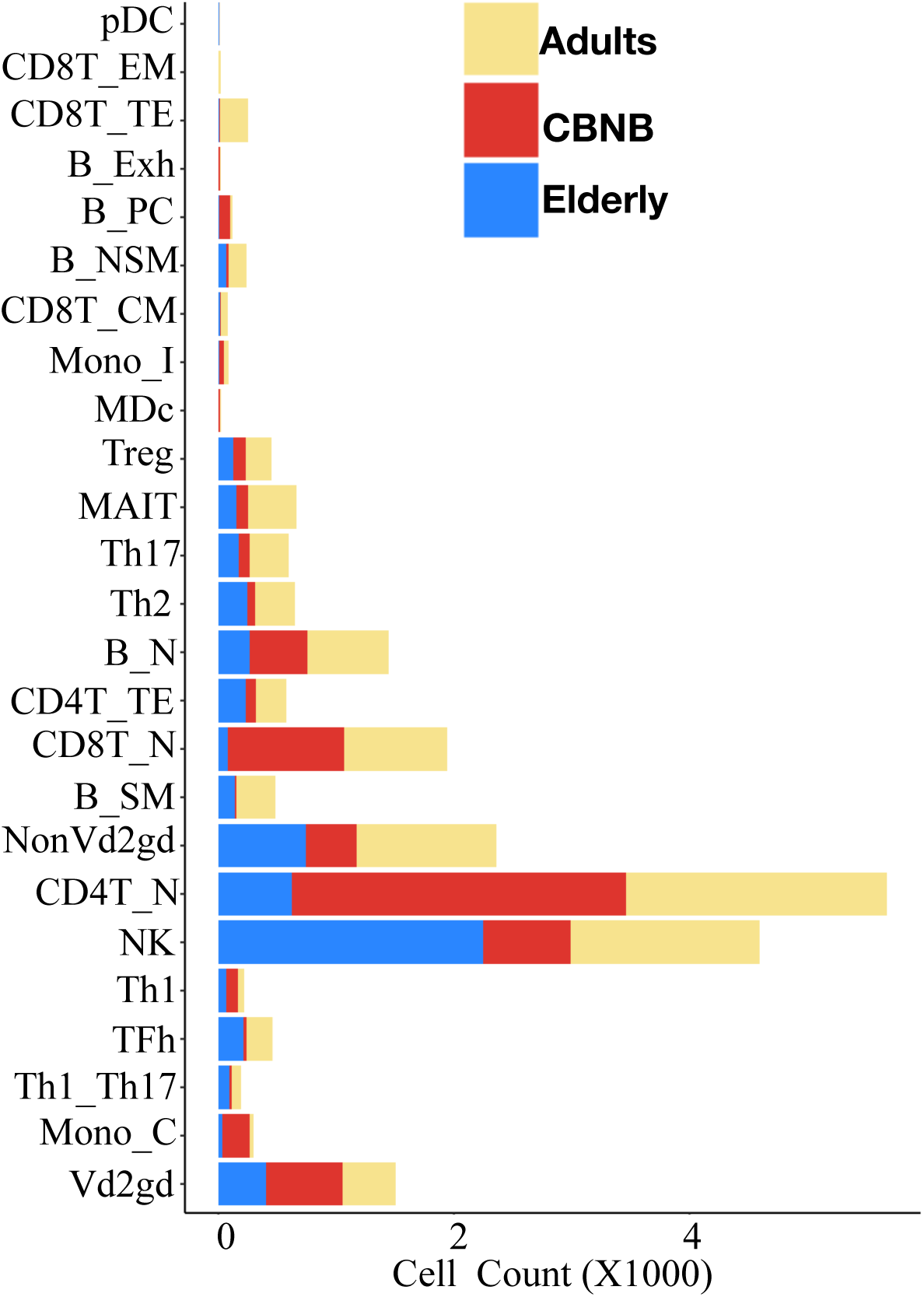
Percentage of immune cell subsets observed in single cell transcriptomics data. Stacked bar-chart shows the cell counts for each subset observed in CBNB, adult and elderly group. Y- axis shows the cell types while X-axis shows cell counts (in thousands).

### KLRB1+ NK cell subset is more central in the elderly group immune cell network

A higher degree of centrality is an indication of more inter-cellular interactions. Accordingly, we hypothesized that the higher degree of centrality will be reflected as higher LR interaction among the subsets. First, we performed t-SNE based dimensionality reduction and clustering for NK cell subsets. Total NK cells (n=4595) from 3 groups clustered into 12 clusters, using lovian clustering algorithm (Fig.5A). Distribution of NK cells on t-SNE plots suggested unique NK cell profile in CBNB, adult, and elderly subjects (Fig.5B). CyTOF data showed, lower frequency (Fig.S4) and degree centrality (Fig.2C, Fig.S2C) for NK cells clusters in CBNB and adult. The two NK subsets (Node87 and Node89) in the elderly immune cell network were unique for its highest degree of centrality and frequency (Fig.5D, Fig. 2C) respectively. These subsets were different in their CD161 expression (Fig.S1A). Node87 expressed CD161 while Node89 was negative for CD161 surface expression. To mirror the CyTOF NK cell phenotype we looked at *KLRB1* (which encodes CD161) gene expression in NK cell clusters from scRNA-seq transcriptomics analysis (Fig.5C). In adults, the majority of cells in clusters 1,4 and 9 were *KLRB1* positive, while in elderly subjects’ *KLRB1* high subsets were in clusters 0 and 2(Fig.5C). We compared the LR interactions between *KLRB1* negative (cluster 3 = NK_3) and *KLRB1* positive (cluster 0 = NK_0) NK cell clusters in elderly subject to mirror the cell phenotype of the Node89 (CD161-) and Node87 (CD161+) subsets from the CyTOF data (Fig.5D). Significantly enriched LR pairs between the NK_0 or NK_3 and all other subsets were obtained using CellChat (18). LR pairs common to both NK_0 and NK_3 were removed and only LR specifically enriched in either NK_0 or NK_3 were counted to understand the interaction potential specific for the *KLRB1* negative and KLRB1 positive NK cell subset. The total LR pair numbers between any subset and NK_0 (*KLRB1*+) were higher than for the NK_3 (*KLRB1*-) subset (Fig.5E). When we analyzed the LR pair directionally, we observed that there were more ligand signals from the NK subset (Fig. S5A) then ligand signal to the NK cell subsets (Fig.S5B).

**Fig. 5.**
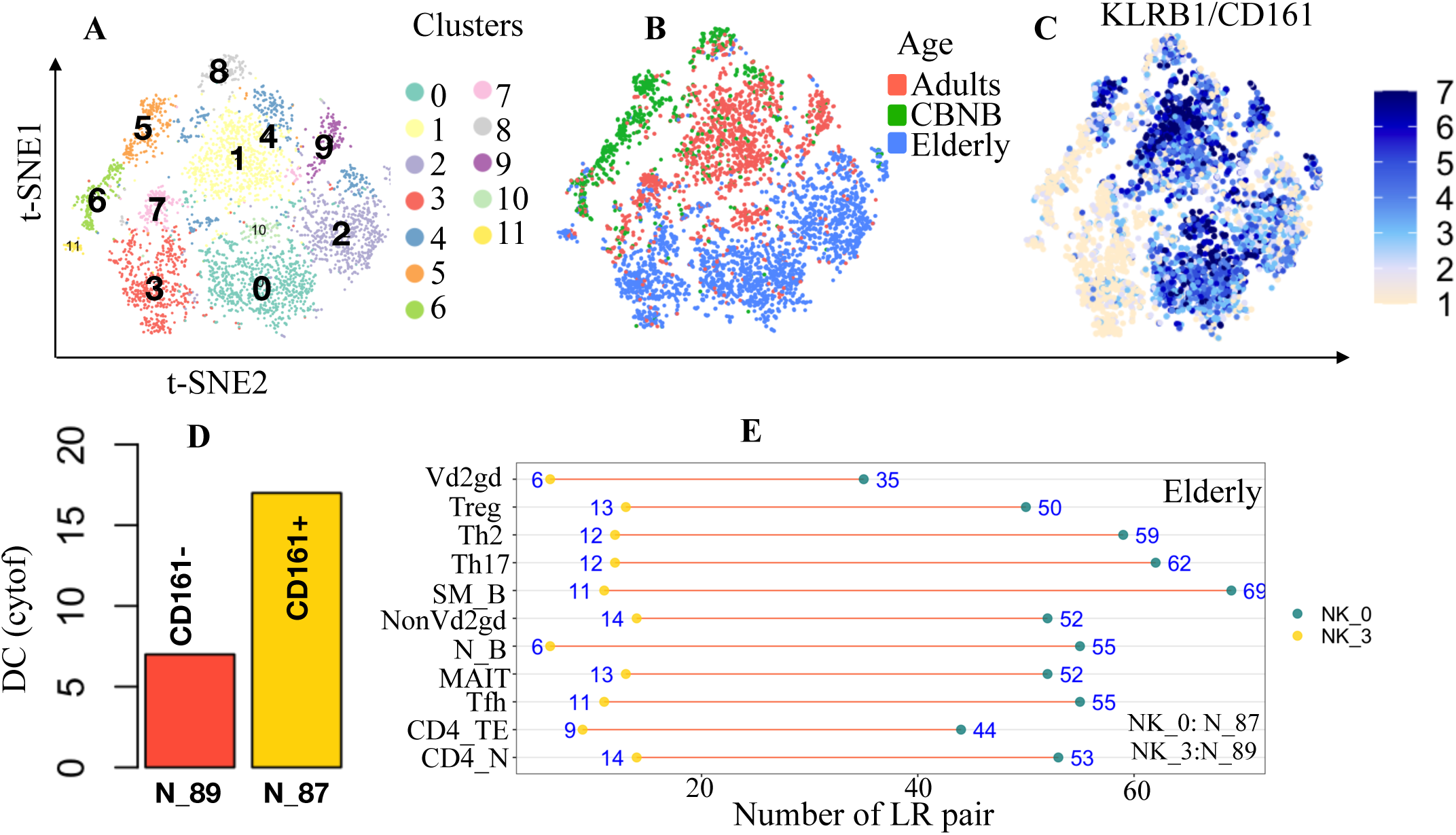
Ligand-Receptor interaction analysis corroborates the high degree centrality of CD161+NK cells. A) Figure shows the clusters of NK cell subsets on the t-SNE plot. B) group information was overlaid on t-SNE plots. C) Expression of KLRB1 gene (KLRB1 genes codes for CD161 protein) is overlaid on t-SNE plot. D) Figure shows the DC of the CD161+ and CD161- NK cell nodes from CyTOF data. E) LR interaction between NK cell cluster NK_0 (KLRB1+) and NK_3 (KLRB1-) similar to CyTOF Node 87 and Node 89 NK cell nodes shown as connected dot chart. The yellow dots represent the number of specific LR pairs enriched between NK_3 and other immune cell subsets shown on Y-axis and green dots denot LR pairs enriched between NK_0 and other immune cell subsets. The line connecting yellow and green dots indicates the difference in interaction potential of these NK cell subsets with other cell subsets.

Hence, higher LR interaction between *KLRB1*+ NK cells with other immune cell subsets corroborates the higher degree centrality and interaction of CD161+ NK cells in the elderly immune cell network.

### Lower LR interactions of CD28- CD8T cells explain exhaustion and immune aging

An increase in CD28- CD8+ senescent and dysfunctional T cells in the elderly population is widely reported(19). CyTOF data in our study recapitulates this observation. CD28-CD8 T cells (Node 69,67) were more abundant than CD28+ CD8T cells in the elderly population, however, CD28- CD8T subset’s degree centrality was lower (Fig.2B). We contrasted LR interactome analysis for CD28- CD8T with CD28+CD8T cells to understand potential differences in inter-subset communication.

To perform LR interaction analysis for CD8 T cells, we pooled all CD8T cells (CD8_ Naive, CD8_EM, CD8_CM, CD8_TE) and reclustered them and performed t-SNE dimensionality reduction for visualisation. We obtained 6 clusters (Fig.6A). Among a total of 2294 CD8 T cells, 1942 were naive CD8 T cells and 253 were terminally exhausted CD8 T cells. Comparatively, central memory(n=21) and effector memory (n=78) T cells were much lower in numbers (Fig.6B).

**Fig. 6.**
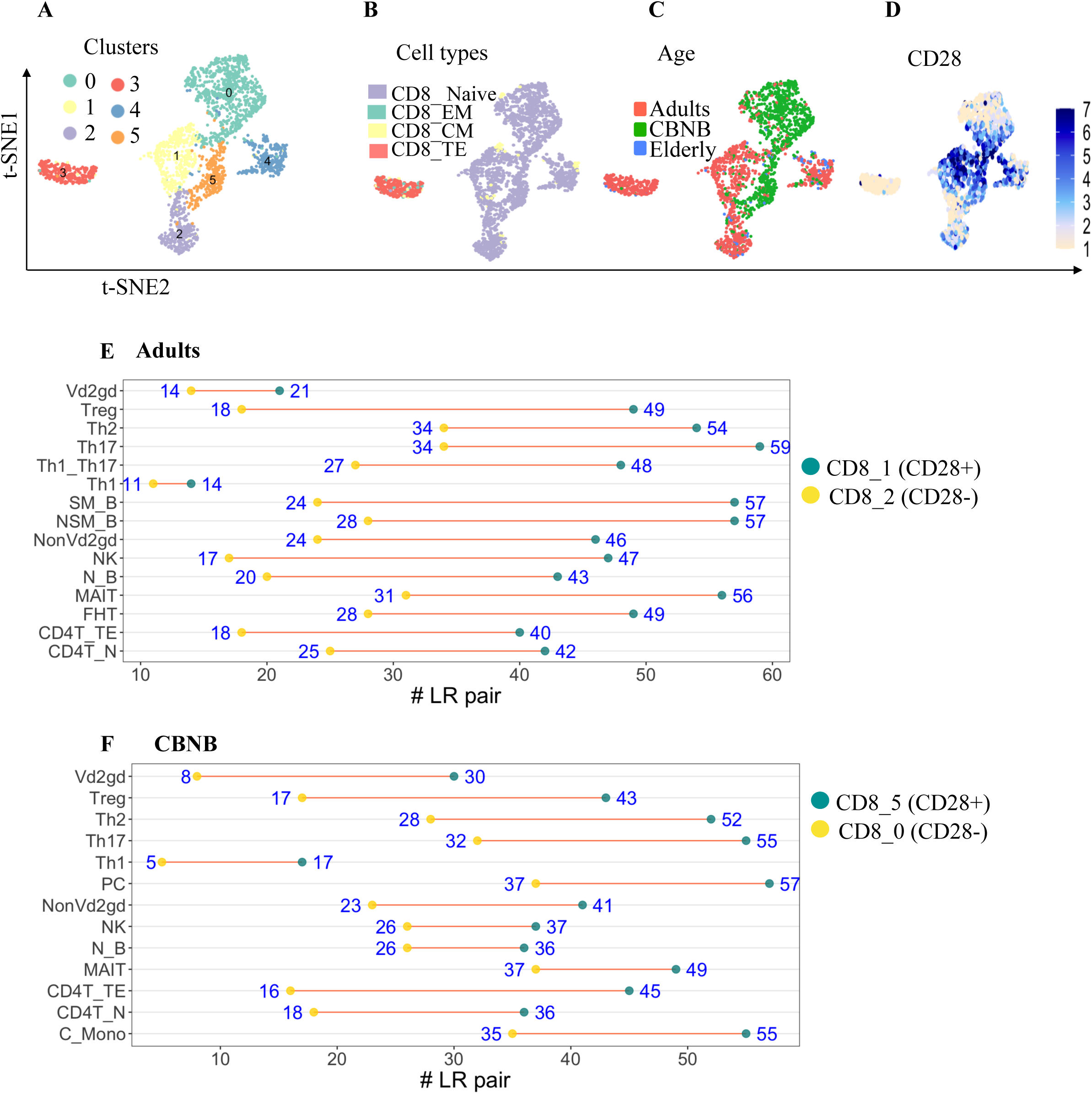
Ligand-Receptor (LR) interaction explains exhaustion phenotype of CD28-CD8 T cells. A) Figure shows the clusters of CD8 T cells derived from scRNA seq overlaid on t-SNE plot. B) Cell type information predicted using MonocoData signature overlaid on t-SNE plot. C) group information was overlaid on t-SNE plots. D) Expression of CD28 gene overlaid on t-SNE plot. E) LR interaction between CD8_2 (CD28-CD8+) and CD8_1 (CD28+CD8+) and all other immune cell subsets from adults shown as connected dot chart. Yellow dots show the number of specific LR pairs enriched between CD8_2 (CD28-) and other immune cell subsets shown on Y-axis and green dots show LR pairs enriched between CD8_1(CD28+) and other immune cell subsets. F) LR interaction between CD8_0 (CD28-CD8+) and CD8_5 (CD28+CD8+) and all other immune cell subsets from CBNB were shown as a connected dot chart. Yellow dots show the number of specific LR pairs enriched between CD8_2/CD8_0 (CD28-) and other immune cell subsets shown on Y-axis and green dots show LR pairs enriched between CD8_1/CD8_5 (CD28+) and other immune cell subsets. The line connecting yellow and green dots indicates difference in interaction potential of these NK cell subsets with other cell subsets.

We tested the LR interactions in *CD28*- and *CD28*+ CD8T cell clusters from adult and CBNB (Fig.6) in scRNA-seq data. In the elderly subject group, we obtained a very low CD8 T cell count (n=100) (Fig.6C). *CD28*+ cluster in adult (CD8_1) was compared with *CD28*- cluster (CD8_2), similarly, CD8_5(*CD28+*) cluster was compared with CD8_0 (*CD28*-) cluster in CBNB. We observed lower LR interaction enriched in *CD28*- CD8 T cells compared to *CD28*+ T cells (Fig.6E, 6F). Lower degree centrality and lower LR interaction of CD28- CD8T cells may thus be one explaintion for exhaustion and loss in communication ability with other immune cell subsets within the immune system.

## Discussion

Immune cells do not function in isolation, but they interact and influence each other and these overall interactions are ultimately reflected as a healthy or diseased phenotype. These cellular interactions lead to a network structure that is specific to any given immune condition and function. Current analysis approaches are mostly reductionist and siloed and fail to represent the complexity of the immune system in health and disease. To the best of our knowledge, no measures were ever used or reported to understand the overall immune cell network in a relational, high dimensionality fashion. Here, we developed a novel systems immunology analytic approach and have created a web application software, CyNET, for high dimensional cytometry data analysis. Our analysis approach nested in CyNET allows an understanding of the immune response holistically at the systems level. We proposed and have validated degree of centrality as an efficient measure of function, which complements the mere calculation of frequency of the immune cell subsets. We developed various systems level measures that capture the relevant biological phenotype. CyNET allows inference of cellular interaction among immune subsets and subnetworks. scRNA sequencing and LR based interactome analysis further validated our inference from the CyNET immune cell network.

We investigated systems level properties such as network density, centralization, and assortativity, to quantify the immune response at the network level(20). We measured these properties in the context of the development of the immune system with age, taking advantage of an already existing, highly curated cytometry dataset (5,11,21). The lifecycle of the immune system starts with the development of a child in the mother’s womb and continues throughout the individual’s life span. Immune cells at birth are immature and start gaining lineage and functional maturity depending on the environmental stimuli. With time, more immune cells gain exhausted characteristics leading to the aging of the immune cells. This immune aging is not only observed in elderly human subjects but is also observed in some autoimmune diseases at early ages. Immature cells during early age will have lower functionality and lower interaction ability with other subsets, a characteristic, which we could capture in our network density analyses. We showed both visually and quantitatively (Fig.1) that network density is lowest in CBNB and highest in adults where cells are fully matured but not as exhausted as in elderly subjects. Network centralization further captures the emergence or change of functional units in the network. Immune cells mature to take up a more specialized function during development or with exposure to antigens and external stimuli. With maturation, these specialized cells dominate in the network in comparison to other subsets. This notion of increased dependence of the network on a few subsets can be quantified using network centralization. As observed, an increase in network centralization with age precisely captures the emergence of more specialized functions. At birth, the majority of the cell subsets are in development and have not undergone enough antigen exposure, thus the network is less centralised.

In contrast, immune cells in the elderly population have been exposed to external stimuli and developed more specialized functions in response to such stimuli. Thus, in elderly populations, some of the immune cells dominate, leading to a more centralised immune cell network. Maturation of the immune cells will lead to preferential attachment of the nodes to generate immune response specific to stimuli. This could be evaluated using assortativity degree of the network. Indeed, increase in centralization score and assortativity degree along the age axis rightly captured the immune cell maturation at the systems level in our study.

Network-level properties can be very useful for monitoring and understanding the system as a whole unit. However, to understand the mechanisms and to optimally manipulate the system for therapeutic development, the understanding of the key nodes within the system is required. Conventionally, in disease vs control comparisons, changes in the frequency of hypothesized immune cell subsets are evaluated to understand the mechanistic relevance of a given subset in health and disease. Here, we propose instead that the centrality of subsets serve as a better measure to understand their functional relevance in patho-physiological systems. For proof of concept, we investigated the change in frequency and degree of centrality of the nodes in the network along the gradient of aging. The overall degree of centrality of naive CD4T cell subsets was lower than other CD4T cell subsets such as Tfh or Treg cells. Thus, higher percentage of naive CD4 T cells in CBNB results in a functionally immature and developing immune system. Tfh frequency in adults and the elderly were similar. However, the degree centrality of Tfh in adults was much higher than in the elderly population. The centrality of Tfh suggested a loss of B cell and antibody secretion functionality in an elderly population. It is well-known and reported that the elderly population is less efficient in generating and maintaining a good antibody titer upon vaccination (22,23). B cells present antigens to Tfh cells and successful engagement of B cells with Tfh protects B cells from death and promotes reentry into the cell cycle. In the germinal center (GC) these B cells mature and exist as antibody producing memory B cells. The loss of Tfh and B cell interactions may further lead to a reduction in B cells that we also observed in an elderly population.

Furthermore, the B cell subsets in the elderly network (Fig.1C) was clustered as a module and were more correlated with each other than with other lineage subsets. Conversely, in adults B cells were more connected with other lineage subsets (Fig.1B). The lower Tfh degree centrality, lower B cell fraction, and isolated module of B cell subsets in the elderly immune cell network accurately highlight the loss of B cell functionality in the elderly population. Like Tfh, the proportion of Th17 (Node 29) was similar in the adult and elderly populations. However, the degree centrality of Th17 subsets was lower in the elderly than in the adult population. Our observation corroborates with a recent study that suggests aging weakens the pathogenicity of Th17 cells (24).

One of the hallmarks of immune aging is the increase in CD28- CD8 T cell frequency(19). CD28 is a costimulatory molecule and is required for successful T-cell activation when the TCR recognizes its cognate antigen presented with MHC. In elderly subjects and autoimmune conditions, constant antigenic exposure and repeated antigenic stimulation lead to loss of CD28 expression on CD8 T cells (19). Low vaccination response in the elderly correlates with a high frequency of CD28- T cells. CD28-CD8T cells negatively impact immune response through various mechanisms. Our study further extends this knowledge and suggests reduced intra-cellular communication potential by altered LR interactions render these cells anergic and dysfunctional in elderly subjects.

NK cell numbers rise with age and the expression of various surface markers on NK cells also changes with age (25–27). We observed a similar increase in NK cell subsets in the elderly population. NK cell expresses many inhibitory and activatory receptors and shows substantial heterogeneity. Among the increased NK cell subsets in elderly group, CD161- subsets were more abundant than CD161+ subsets (Fig. 2C). Only one NK cell subset (Node 87) was positive for CD161 expression (Fig.S1A). CD161 expression on NK cells marks responsiveness to pro-inflammatory cytokines, increased expression of integrins, and tissue residency markers (28). CD161 expression is lost in CMV-induced terminally differentiated NK cells (28). Human CMV-induced adaptive NK cell subset also shows downregulation of CD161 gene expression, and these adaptive NK cells are suggested to have emerged for immunosurveillance of infected cells with improved proliferation and survival capacity in inflammatory conditions (29). The lower degree centrality, and LR interaction potential of the CD161- NK cell subsets compared to CD161+ NK cell subsets corroborates and explains the reduction in signaling proteins and response to proinflammatory cytokines by CD161- NK cells. CD161+ NK cells are inhibitory and probably enable immune homeostasis and relative contraction of these cytokine-responsive subsets in the elderly population may further add to immune aging.

Altogether, we have broadened our high dimensionality approach to the analysis of the Immunome across the age gradient with an in-depth understanding of the relational networks, which shape mechanisms of immune responses and their evolution. Some of our findings show a surprising discordance between frequency and centrality for given, mechanistically relevant, immune subsets, thus underscoring the importance of integrating the traditional approach based on cell frequency in a given subset with network analyses capable of defining the relational relevance of such subsets. We offer here to the community (https://epicimmuneatlas.org) the necessary tools and dataset to apply this sort of analysis to various conditions, in the hopes of expanding and strengthening this approach.

## Methods

### CyNET web application

The CyNET web application was created using the R shiny web framework. For network analysis igraph R package was used, and for network visualisation visNetwork, and ggplot2 R packages were used.

### CyTOF data analysis

CyTOF data for CBNB, adult and elderly subjects were downloaded from EPIC (epicimmuneatlas.org). Data was clustered into 100 clusters for both Panel 1 and Panel 2 dataset. Cluster frequency and median marker expression data was downloaded and used for network analysis. T cells and NK cells from Panel 1 data, B cells and Monocyte/DC data from Panel 2 data set was merged together create a full lineage set data matrix. To balance the sample number in each group 17 sample (Maximum number of samples available for elderly group in the dataset) from each category was randomly taken for correlation network analysis. To create the correlation network correlation (R2 = 0.5) and p-value (<0.05) cutoff was applied to establish a n edge between the nodes. The immune cell clusters were the nodes of the network and the edges between them was considered potential direct or indirect interaction. To create the network and calculate the network properties igraph R package and custom R code was used. To visualise the network visNetwork R package was used.

### Healthy Human blood sample collection

Peripheral blood samples were collected from CBNB(n=5), adult(n=6), and elderly(n=4) healthy human subjects. The subject’s demographic information is provided in Supplementary Table S1. The paediatric and adult samples were obtained with informed consent from KK Women’s and Children’s Hospital, Singapore, with ethical approval from the SingHealth Centralized Institutional Review Board. The elderly samples were obtained from the National University Hospital System, Singapore.

### Peripheral blood mononuclear cells (PBMCs) preparation

Peripheral blood samples were collected in ethylene diamine tetra acetic acid tubes (EDTA). PBMCs were isolated from blood samples using Ficoll-Paque (GE Healthcare) gradient centrifugation within 6 hrs from recruitment, and were stored in liquid nitrogen until further use.

### Single cell Library preparation and sequencing

PBMCs were thawed and rested for 30 minutes. Cells were washed with PBS twice and Single cells were encapsulated into droplets with a gel bead in the emulsion (GEM) method using the 10x Chromium controller. The Chromium Single Cell 3ʹ gene expression protocol (version 3 chemistry,10x Genomics) was used. Downstream library construction was performed using the 10x 3ʹ Gene Expression Library Construction kit, and barcoding was carried out with i7 Illumina adaptor indexes. Pooled libraries were then sequenced on the Illumina HiSeq 4000 platform using paired-end 151-bp reads to achieve 50,000 reads per cell for gene expression

### Single cell data analysis

Raw reads for transcriptomic analyses were aligned to the human genome (hg19- 3.0.0) using Cell Ranger version 7 software. CellRanger’s count utility was used to count the features. Cell barcodes and feature count matrices were created by aggregating filtered feature counts of each sample using the Cell Ranger aggregate utility. Subsequent data normalization and analysis were performed using the Seurat R package and custom R code. Cell count data were quality controlled and filtered based on cellular complexity (number of genes per cell) and mitochondrial reads. Cells with between 300 and 5,000 genes and mitochondrial percent reads less than 20 were kept for analysis. Data scaling, normalization, variable gene identification and clustering were performed using the Seurat pipeline. SCT transform method from Seurat R package was used for batch normalisation. Batch normalised data was used for imputation of gene counts using ALRA algorithm (16). Imputed gene count matrix was further used for cell type annotation using SingleR package and MonacoData set signature. For clustering within the annotated subsets Lovian clustering was performed.

### Ligand Receptor analysis

To find the potential interaction between immune cell subsets/clusters, networks of clusters created based on enriched LR pairs between the clusters. CellChat package (18) was used to find significantly enriched LR pairs between the immune subsets. For visualisation and analysis visNetwork, ggplot2 and customs R code was used.

## Data availability

The software application can be accessed on https://epicimmuneatlas.org.

Raw RNA sequencing data are from human subjects and will be provided upon request in alignment with institutional and grantor guidelines.

## Code availability

Custom code will be given on reasonable request.

## Acknowledgement

We thank all participants and their families who gave consented to the use of their biological samples for this study. This study was supported by grants from the NMRC (MOHIAFCAT2/005/2015, NMRC/OFLCG/002/2018, CIRG19may0052), MOH-STaR19nov-0002, A*STAR PEC21-H22P0M0003, Duke-NUS and SingHealth AMC core funding is gratefully acknowledged. This research is also supported by the Singapore Ministry of Health’s National Medical Research Council under its Centre Grant Programme (MOH-000988). We thank Associate Professor Ng Tze Pin, National University Hospital System, Singapore, for contributing the elderly samples to the study.

## Author contributions

P.K performed the experiments and bioinformatics analysis. S.L.P, S.N.H, N.B.S, V.R.C, W.X.G.A, J.Y.L performed the experiments. J.G.Y, Y.K.T and T.A. recruited the human subjects. T.A, J.G.Y, L.R.K., W.X.G.A, M.W. participated in study design and manuscript preparation. P.K, and S.A conceived and lead the study and wrote the manuscript. SA arranged funding for the study

## Extended data figure legends

**Fig. S1.**
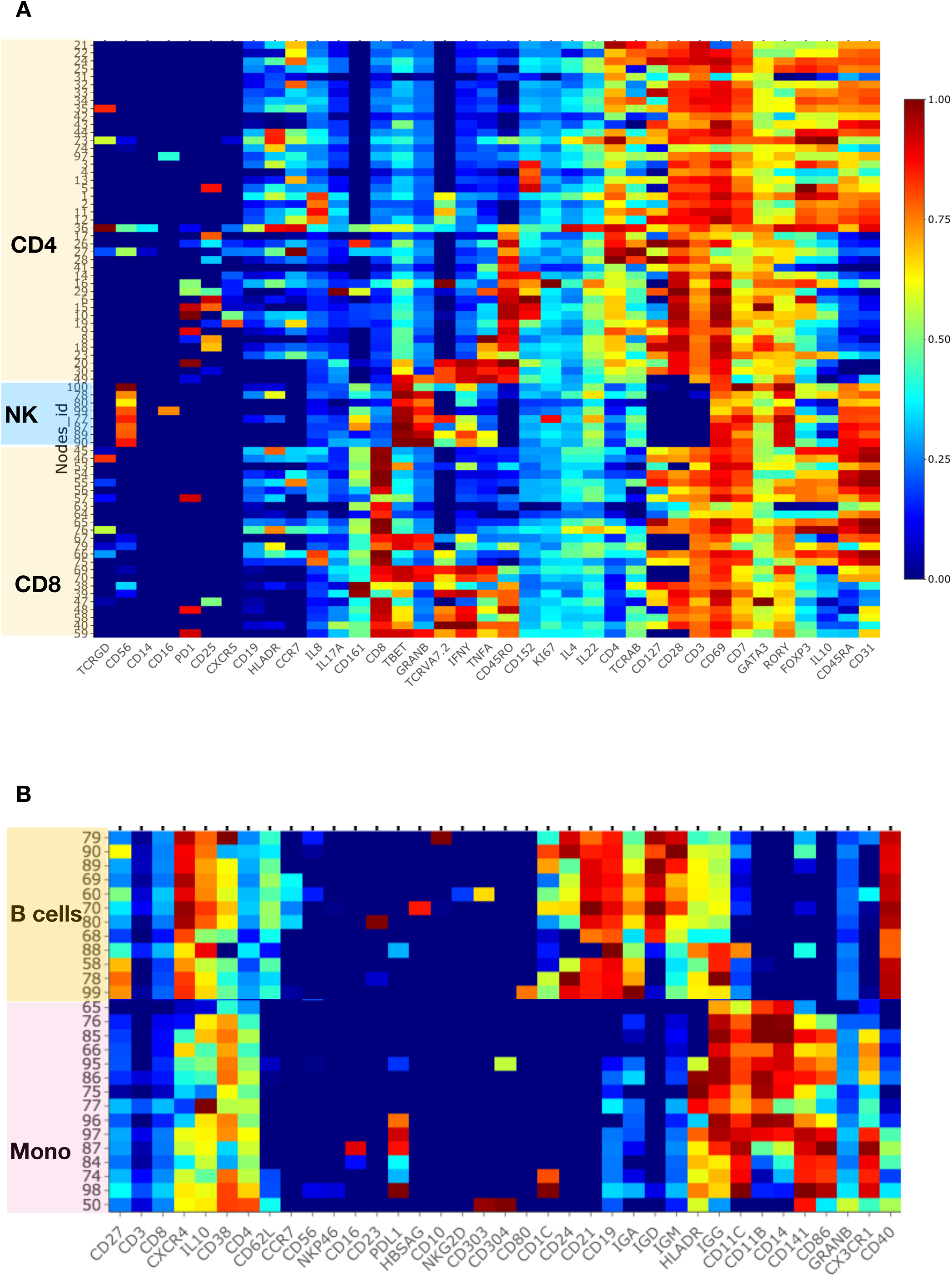
Phenotype of nodes (Immune cell subsets) Normalised median antibody expression for all the nodes from A) Panel 1, and B) Panel 2 plotted as heatmap

**Fig. S2.**
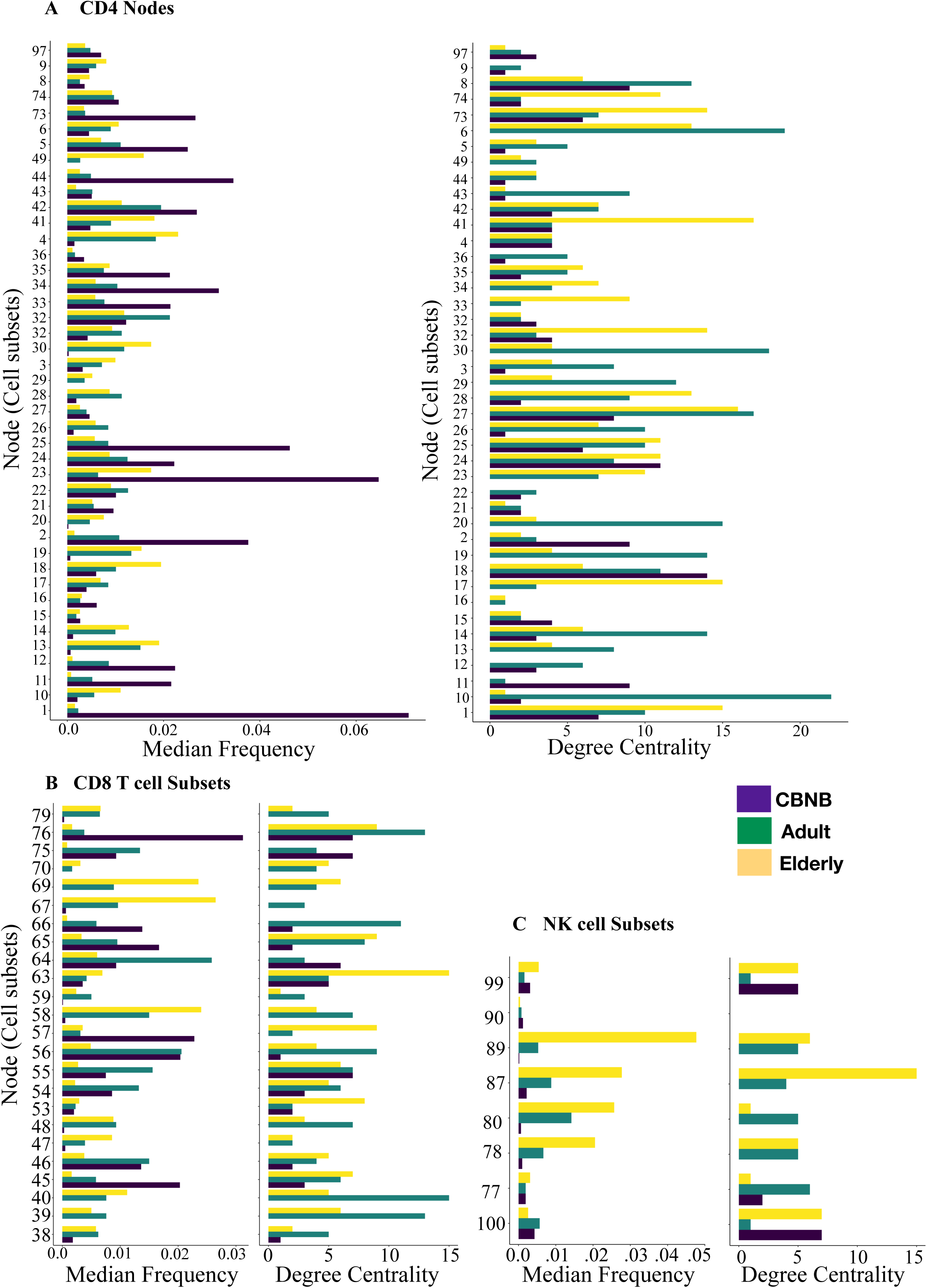
Degree Centrality (DC) and frequency of immune cell subsets from panel 1 dataset. Median frequency and DC of each node in A) CD4 T cell subsets C) CD8 T cell subset D) NK cell subsets were plotted as bar-plot. Colour of the bars in bar-plot shows the human sample group information. Each node number is shown on Y -axis while median frequency and DC was shown on X-axis.

**Fig. S3.**
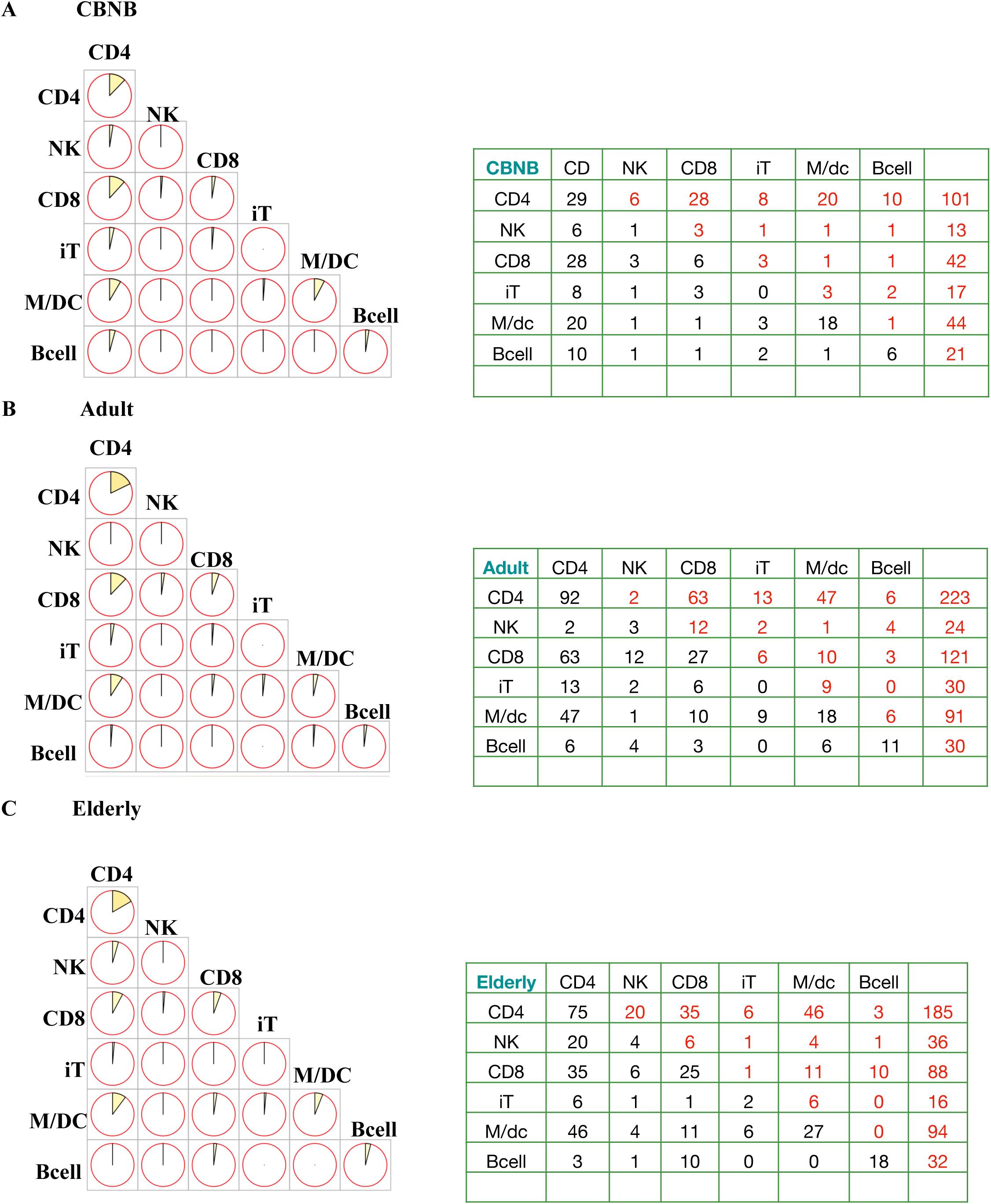
Interaction between the major lineage cell subsets. Total number of edges between major cell lineage subsets in network of A) CBNB, B) Adult, C) Elderly were shown as a pie-chart and the number of edges between the subsets were listed in a table. Size of the pie in the circle reflects the fraction of edges normalised for total number of edges respective each network.

**Fig. S4.**
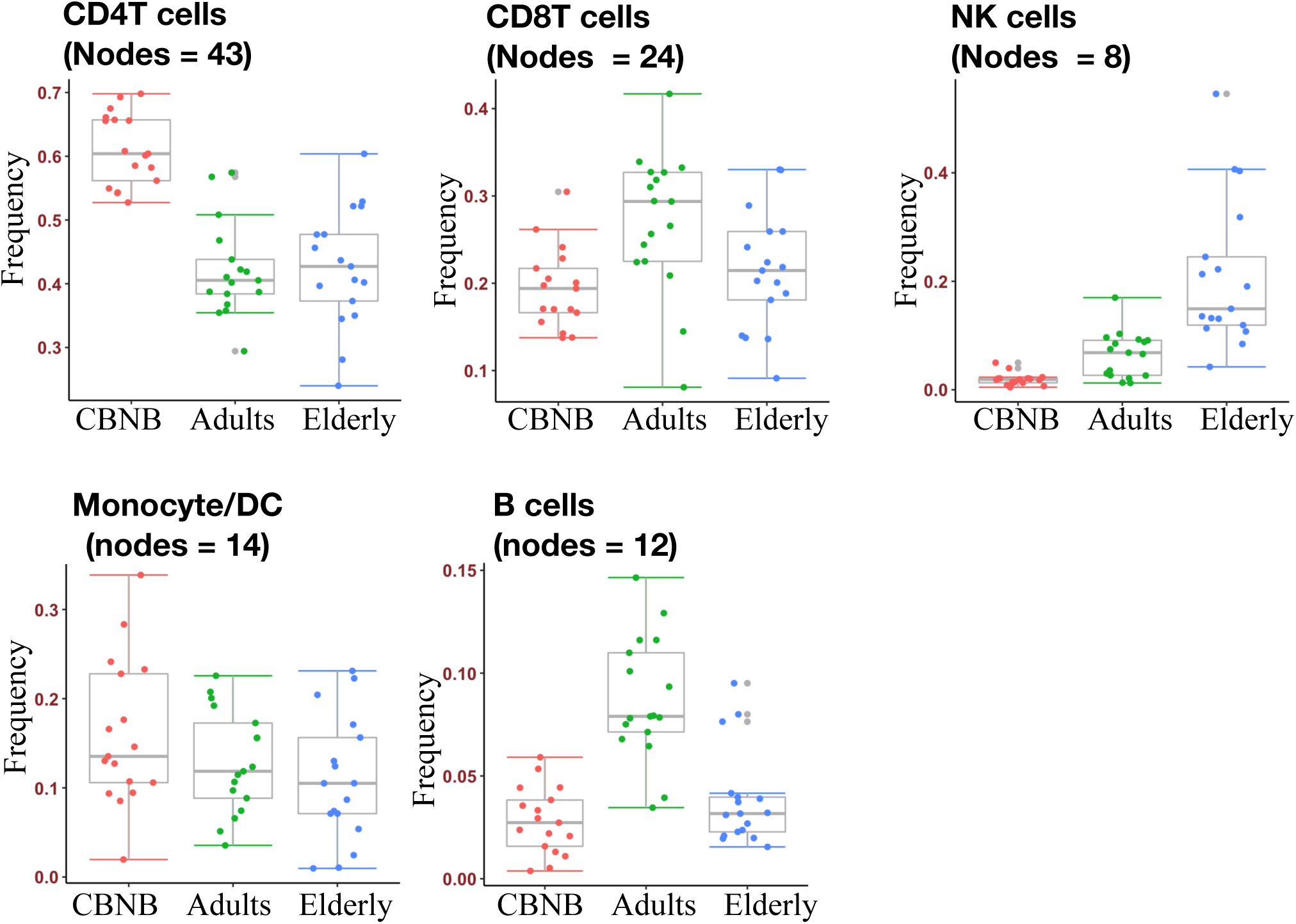
Frequency of Major cell lineage in from CyTOF data. Box and whisker plot showing frequencies of the major cell lineages (CD4 T cells, CD8 T cells, NK cells, Monocytes/DCs, B cells) in CBNB, adult and elderly groups.

**Fig. S5.**
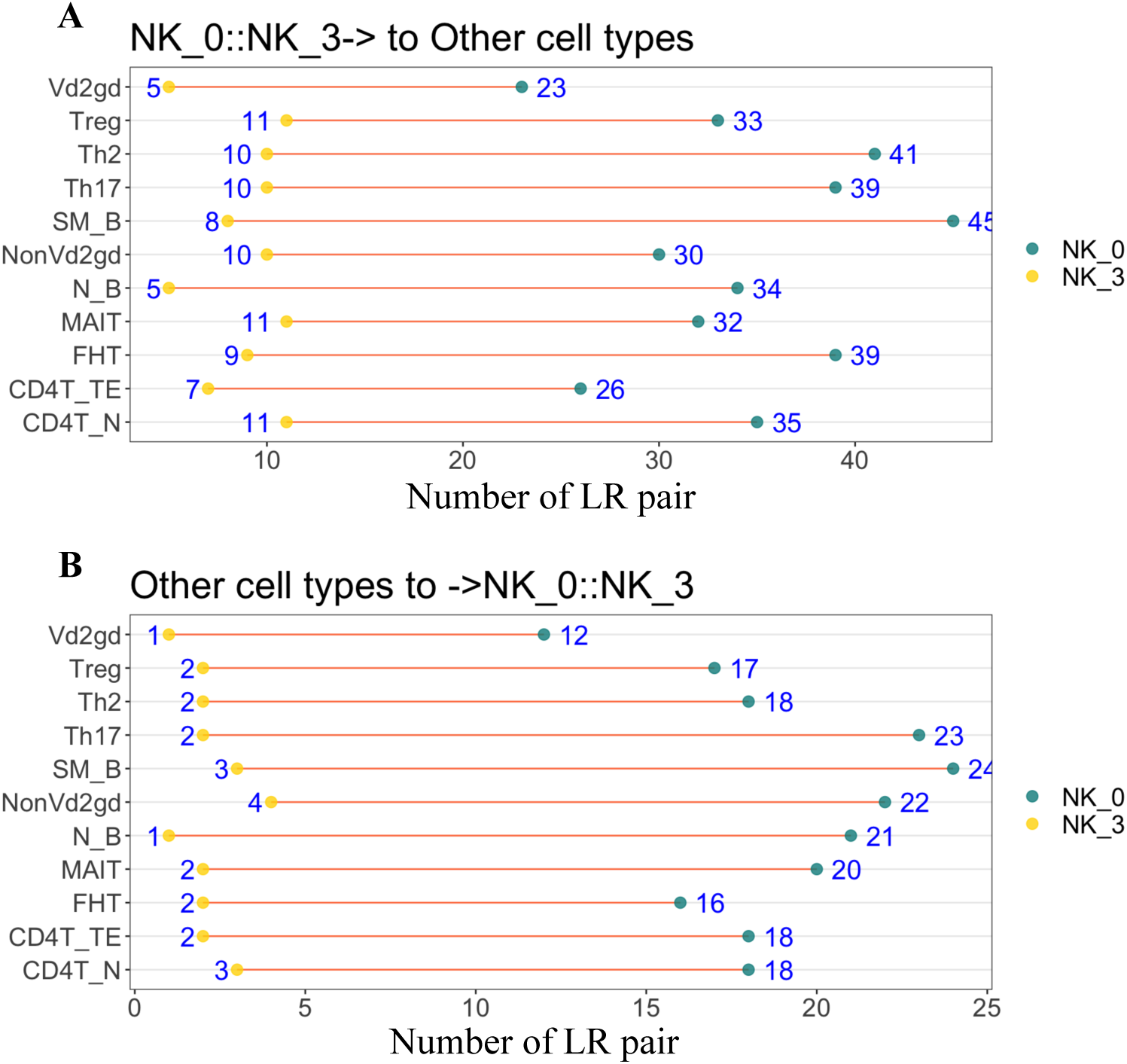
Directionality of Ligand-Receptor interaction analysis in NK cells. Ligand-Receptor (LR) interaction analysis shown separately for ligands signalling from A) NK cell subsets to other immune cell subsets or B) ligands signalling from other subsets to NK cell subsets. LR interactions between NK cell cluster NK_0 (KLRB1+) and NK_3 (KLRB1-) and other cell subsets were shown as a connected dot chart. Yellow dots show the number of specific LR pairs enriched between NK_3 and other immune cell subsets shown on Y-axis and green dots show LR pairs enriched between NK_0 and other immune cell subsets. Lines connecting yellow and green dots indicates the differences in interaction potential of these NK cell subsets with other cell subsets.

## Supplementary Information

Supplementary Table S1: Human Subject demographic information

Supplementary Table S2: Cells counts of identified cell subsets in scRNA-seq data

Supplementary Table S3: Differential genes expressed in naive CD4T cells (CD4TN) compared to resting CD4 T cells (restCD4T)

